# Decoding fMRI Data: A Comparison Between Support Vector Machines and Deep Neural Networks

**DOI:** 10.1101/2023.05.30.542882

**Authors:** Yun Liang, Ke Bo, Sreenivasan Meyyappan, Mingzhou Ding

## Abstract

Multivoxel pattern analysis (MVPA) examines the differences in fMRI activation patterns associated with different cognitive conditions and provides information not possible with the conventional univariate analysis. Support vector machines (SVMs) are the predominant machine learning method in MVPA. SVMs are intuitive and easy to apply. The limitation is that it is a linear method and mainly suitable for analyzing data that are linearly separable. Convolutional neural networks (CNNs), a class of AI models originally developed for object recognition, are known to have the ability to approximate nonlinear relationships. CNNs are rapidly becoming an alternative to SVMs. The purpose of this study is to compare the two methods when they are applied to the same datasets. Two datasets were considered: (1) fMRI data collected from participants during a cued visual spatial attention task (the attention dataset) and (2) fMRI data collected from participants viewing natural images containing varying degrees of affective content (the emotion dataset). We found that (1) both SVM and CNN are able to achieve above chance level decoding accuracies for attention control and emotion processing in both the primary visual cortex and the whole brain with, (2) the CNN decoding accuracies are consistently higher than that of the SVM, (3) the SVM and CNN decoding accuracies are generally not correlated with each other, and (4) the heatmaps derived from SVM and CNN are not significantly overlapping. These results suggest that (1) there are both linearly separable features and nonlinearly separable features in fMRI data that distinguish cognitive conditions and (2) applying both SVM and CNN to the same data may yield a more comprehensive understanding of neuroimaging data.

**Key points:**

- We compared the performance and characteristics of SVM and CNN, two major methods in MVPA analysis of neuroimaging data, by applying them to the same two fMRI datasets.
- Both SVM and CNN achieved decoding accuracies above chance level for both datasets in the chosen ROIs and the CNN decoding accuracies were consistently higher than those of SVM.
- The heatmaps derived from SVM and CNN, which assess the contribution of voxels or brain regions to MVPA decoding performance, showed no significant overlap, providing evidence that the two methods depend on distinct brain activity patterns for decoding cognitive conditions.

## Introduction

Functional magnetic resonance imaging (fMRI) exploits blood-oxygen-level-dependent (BOLD) contrasts to map neural activities (Mahmoudi et al., 2012). Conventional analyses of fMRI data using methods such as the general linear model (GLM) compare BOLD activities evoked by different experimental conditions at the single voxel level or at the level of a region of interest (ROI) to gain insights into the neural basis of cognitive functions. These conventional analyses, referred to as the univariate approach here, have yielded much of our current understanding of the neural underpinnings of human cognition. Like any methods, the univariate approach to fMRI data analysis has limitations. For example, for a voxel to be considered activated by a cognitive event, it has to be consistently activated in a population of participants, which is often difficult to achieve given the idiosyncratic nature of neural responses to subtle experimental manipulations and can thus lead to failure to detect the presence of neural signals (e.g., (Haxby et al., 2011)). At the ROI level, averaging across voxels disregards the information contained in the heterogeneous response patterns, again reducing the sensitivity of our ability to detect differential neural responses to different experimental conditions.

Multivoxel pattern analysis (MVPA), which can be performed at the single subject level and takes into account the spatial variations of the BOLD activity across voxels, overcomes the limitations of the univariate approach and have become the main approach for providing information that complements the univariate approach (Haynes, 2015; Lewis-Peacock & Norman, 2014; Mitchell et al., 2003; Norman et al., 2006; Tagliazucchi & Laufs, 2014; Tagliazucchi et al., 2012). Currently, the support vector machine (SVM) (Sain, 1996) is the most widely used MVPA method for analyzing fMRI data (Bonnici et al., 2013; LaConte et al., 2005; Song et al., 2011). When decoding between different cognitive conditions using SVM, a classifier is trained on training data and tested on testing data, and above-chance level decoding accuracy is taken as evidence of the involvement of the ROI in the cognitive function being tested. The MVPA literature has grown significantly in recent years (Baucom et al., 2012; Kotz et al., 2013; Said et al., 2010; Sitaram et al., 2011). Novel insights not possible with traditional univariate methods have emerged in all fields of cognitive neuroscience. We used the MVPA approach to decode neural responses to natural images containing varying degrees of emotional content and reveal the existence of affect-specific neural representation in the retinotopic visual cortex (Bo et al., 2021). Besides decoding accuracy, heatmaps are another SVM construct that can be derived from the classifier, which provide information on the contribution of different voxels to the classifier performance. To date heatmaps are far less utilized than decoding accuracy. We have used SVM heatmaps to uncover the attentional control microstructures in the dorsal attention network (DAN) (Rajan et al., 2021). Despite SVM’s success, the linear nature of the method is both a strength and a weakness. Being linear, SVM is intuitive to understand and computationally efficient. On the other hand, being linear, SVM is effective only when the data is linearly separable. Recent studies suggest that the mapping between neural activity and cognitive functions may be nonlinear (Birn et al., 2001) and as such linear SVM may not be able to effectively characterize the underlying brain patterns (Farahani et al., 2019). Another known weakness of SVM is its limited ability for handling high-dimensional data; the need for expert feature selection to reduce data dimensionality may bias the results (Vieira et al., 2017).

The advent of AI-inspired methods such as deep neural networks (DNNs) has the potential to overcome SVM’s limitations and provide information that complements the SVM. DNNs, especially the Convolutional Neural Networks (CNNs), have emerged as a technique for extracting effective features from multivariate neuroimaging data (Abrol et al., 2021). When used to decode different experimental conditions, a CNN model is trained on training data and tested on testing data, which is similar to SVM. Both decoding accuracy and heatmaps can be derived from the trained CNN models. Recently, we applied a CNN model to fMRI data recorded from patients suffering from trigeminal neuralgia to reveal novel insights into the generation and perception of TN pain, which were not possible with other methods (Liang et al., 2022). Given the prevalence of SVM and the emerging significance of CNN in MVPA analysis of neuroimaging data, we consider it timely to compare CNNs and SVMs by applying them to the same datasets. For this purpose, two well-characterized datasets were considered: (1) fMRI data recorded from participants performing a cued spatial attentional task (attention dataset) and (2) fMRI data recorded from participants viewing natural images containing varying levels of affective content (emotion dataset). The main quantities of interest for comparison are the decoding accuracy and the heatmap. Two hypotheses were tested: (1) CNN will achieve higher decoding accuracy than SVM and (2) CNN and SVM rely on different features for decoding and classification.

## Materials and Methods

Two fMRI datasets were used to compare SVM and CNN decoding. To facilitate the comparison, recording, preprocessing, and analyses were carried out identically for both datasets. We note that these datasets have been used in previous publications to address different questions (Bo et al., 2021; Meyyappan et al., 2021).

### The Attention Dataset

The experimental protocol was approved by the Institutional Review Board of the University of Florida. Twenty right-handed healthy volunteers (15 men and 5 women; mean age: 24.65 ± 2.87 years) with normal or corrected-to-normal vision and no history of neurological or psychological disorders provided written informed consent and participated in the study. This dataset has been used in prior publications to address different questions (Meyyappan et al., 2021).

The participants performed a cued visual spatial/feature attention experiment while simultaneous EEG-fMRI were recorded (only the fMRI data were considered here). The start of each trial was signaled by an auditory cue, which directed the participants to direct their attention to either a spatial location (left or right) or a color (red or green). Following a delay period, varied randomly from 3000 to 6600 ms, two colored rectangles (red or green) were presented for a duration of 200 ms, with one in each of the two peripheral locations. The subject’s task was to report the orientation of the rectangle (target) appearing at the cued location or having the cued color. For feature (color) trials, the two rectangles displayed were always of the opposite color; for spatial trials, the two rectangles were either of the same color or the opposite color, see **Figure 1A** (Meyyappan et al., 2021). For this study, we mainly focused on the cue-related activity, reflecting preparatory or anticipatory attention, in the spatial attention conditions (Meyyappan et al., 2021); the feature attention conditions were not considered.

**Figure 1:**
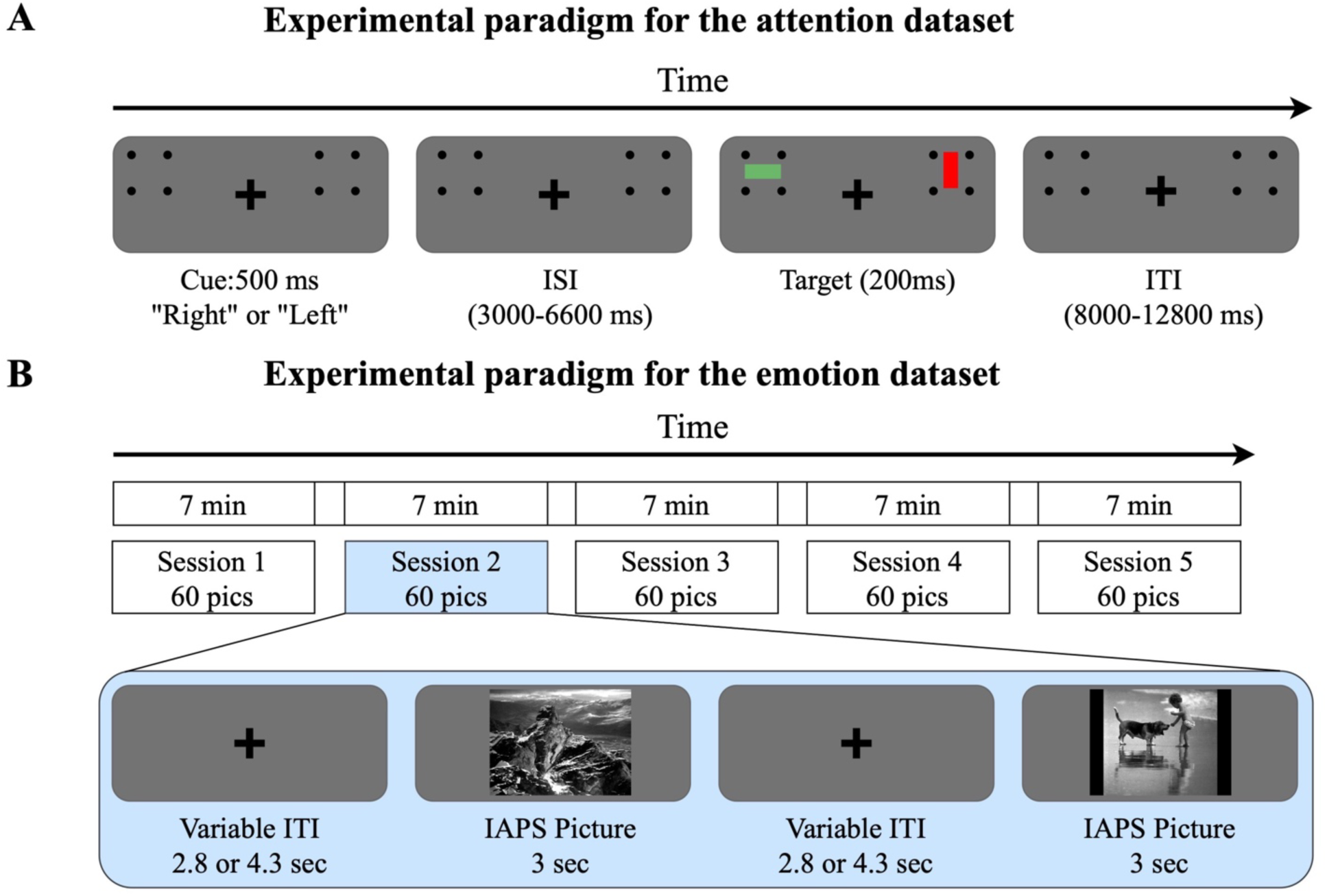
Experimental paradigms. (A) Attention dataset. Each trial begins with an auditory cue lasting 500 ms, instructing the subject to covertly attend to a spatial location (left or right). After a variable cue-to-target delay ranging from 3000 to 6600 ms, two colored rectangles are displayed for 200 ms, one in each of the two peripheral locations. Participants are then asked to report the orientation of the rectangle (horizontal or vertical) appearing in the cued location. An intertrial interval (ITI) follows, randomly varied from 8000 to 12,800 ms after the target onset, before the start of the next trial. (B) Emotion dataset. The experiment consisted of five sessions, each lasting 7 minutes. In each session, a total of 60 International Affective Picture System (IAPS) pictures, including 20 pleasant, 20 unpleasant, and 20 neutral, were presented in random order. Each picture was displayed for 3 seconds, followed by a fixation period referred to as inter-trial interval (ITI) lasting 2.8 or 4.3 seconds. Participants were required to fixate on the cross in the center of the screen throughout the session.

### The Emotion Dataset

The experimental protocol was approved by the Institutional Review Board of the University of Florida. A total of 26 healthy volunteers with normal or corrected-to-normal vision gave written informed consent and participated in the study. Two participants withdrew from the experiment. Four additional participants were discarded due to excessive movements inside the scanner. The data from the remaining 20 participants were analyzed and reported here (10 men and 10 women; mean age: 20.4 ± 3.1 years) (Bo et al., 2021).

The participants viewed 60 gray-scaled pictures including 20 pleasant, 20 unpleasant, and 20 neutral pictures from the International Affective Picture System (IAPS) library while simultaneous EEG-fMRI was recorded (only the fMRI data were considered here (Lang et al., 1997)). There were 5 runs. The 60 pictures were presented in random order in each run, see **Figure 1B**. The participants passively viewed the images in the scanner. No response was required from them. For this study, we mainly focused on the unpleasant pictures and neutral pictures.

### fMRI acquisition and preprocessing

For both datasets, the functional MRI data were collected on a 3T Philips Achieva scanner (Philips Medical Systems), with the following parameters: echo time, 30 ms; repetition time, 1.98 s; flip angle, 80◦; slice number, 36; field of view (FOV), 224 mm; voxel size, 3.5 × 3.5 × 3.5 mm; matrix size, 64×64. Preprocessing was carried out using the statistical parametric mapping toolbox (SPM) and custom scripts written in MATLAB, including the following steps: slice timing correction, realignment, spatial normalization, and smoothing. Slice timing correction was conducted using sinc interpolation to correct for differences in slice acquisition time within an EPI volume. The images were then spatially realigned to the first image of each session by a 6-parameter rigid-body spatial transformation to account for head movement during data acquisition. Each participant’s images were then normalized and registered to the MNI space. All images were further resampled to a voxel size of 3 × 3 × 3 mm and spatially smoothed using a Gaussian kernel with 7 mm FWHM. The preprocessed signal was further passed through a high-pass filter with a cutoff frequency set at 1/128 Hz to attenuate lower frequency noise.

### Regions of Interest (ROIs)

For both datasets, the stimuli are visual, and as such the primary visual cortex (V1) plays an essential role in extracting and processing basic visual features of the stimuli. It is also known that V1 is subject to influence and modulation of higher-order visual structures depending on the task conditions. We first performed the comparison between SVM and CNN in primary visual cortex (V1). The goal was to examine whether SVM and CNN exhibit comparable classification performance and make classifications of cognitive states based on similar visual features. V1v (V1 ventral) and V1d (V1 dorsal) in a recently published probabilistic visual retinotopic atlas (Wang et al., 2015) were combined to form the V1 ROI (**Figure 2A**). We next performed the comparison between SVM and CNN at the whole brain level because cognitive functions such as attention and emotion engage large-scale distributed brain networks. The goal was to examine whether SVM and CNN exhibit similar classification performance and make classifications of cognitive states based on similar brain regions. The 129 brain regions in the Lausanne Brain Atlas are combined to form the whole brain ROI (Daducci et al., 2012); see **Figure 2B**.

**Figure 2:**
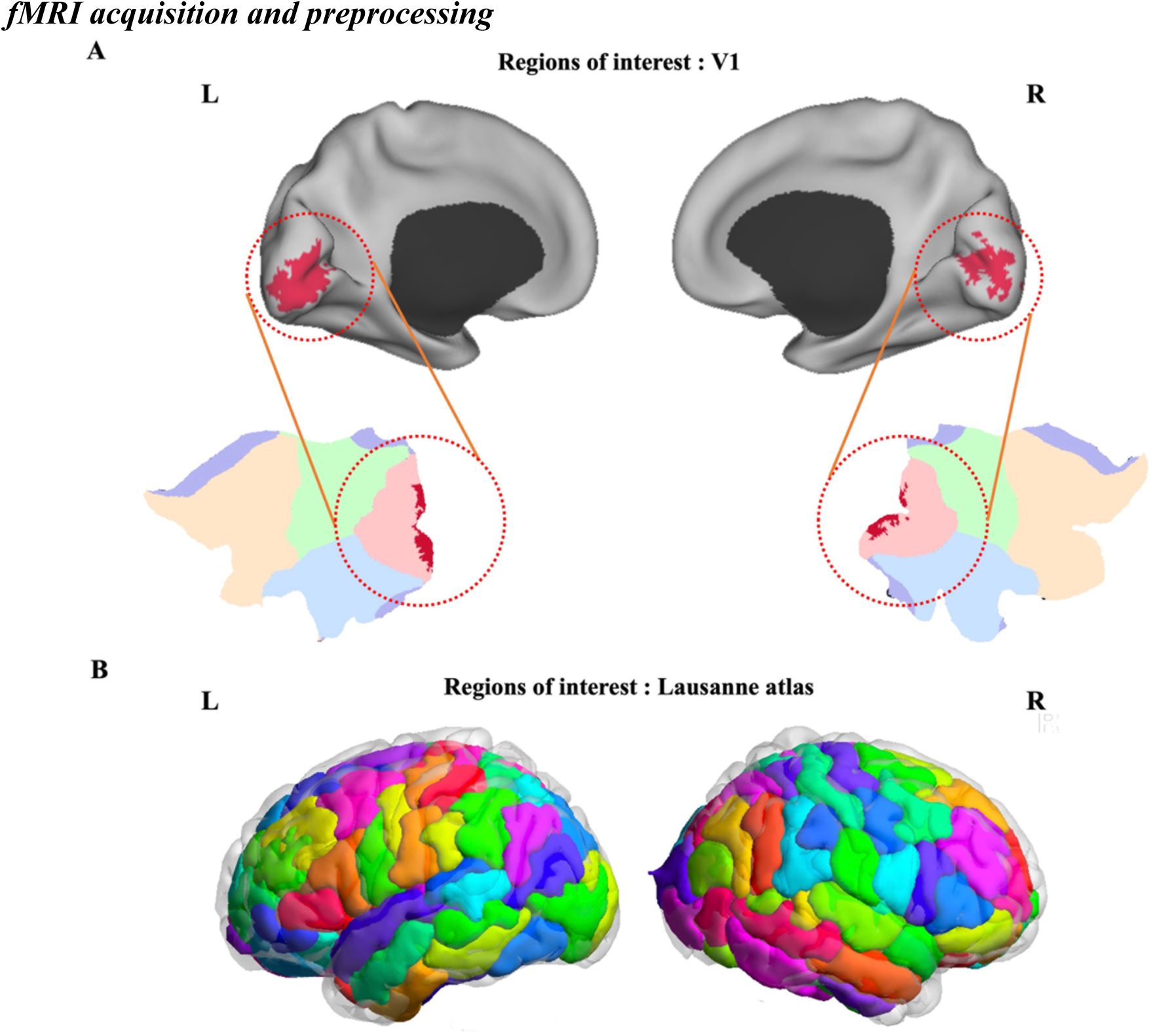
Regions of interest. (A) V1 ROI. (B) Whole brain ROI based on Lausanne Atlas (129 regions).

### Univariate Analysis with GLM

As a starting point, we first analyzed the data using the univariate approach, in which we adopted the standard GLM method, as implemented in the SPM toolbox. For the attention dataset, eight task-related events were included as regressors in the GLM model, including five regressors to model the cue-related BOLD activity, two regressors to account for BOLD responses evoked by validly and invalidly cued targets separately, and one regressor to model trials with incorrect responses. For the emotion dataset, four task-related events were included as regressors in the GLM analysis, including three stimuli-related regressors and one fixation regressor. For both datasets, the Hemodynamic Response Function (HRF) used in the GLM analysis was the default HRF provided by the SPM toolbox, with a delay of 6 seconds. At the group level, fMRI activation maps were obtained by applying a parametric one-sample t-test and thresholded at a significance level of p<0.05 after correcting for multiple comparisons using the false discovery rate (FDR) method.

### Single-Trial Estimation of fMRI-BOLD

Since MVPA is performed at the single-trial level, we applied a beta series regression method to estimate the BOLD response on each trial in every voxel for both attention and emotion datasets (Mumford et al., 2012). Specifically, for both datasets, the trial of interest was represented by one regressor, and all the other trials were represented by another regressor. Six motion regressors were also included to account for any movement-related artifacts during the scan. The process was repeated for all trials. The single-trial beta responses were fed into SVM and CNN for decoding analysis and heatmap generation.

### MVPA: SVM

SVM as implemented in the LibSVM (Chang, 2011) was used to decode cue left vs. right (spatial attention control) for the attention dataset using both the V1 ROI and the whole brain. For the emotion dataset, we decoded the brain patterns evoked by unpleasant vs. neutral stimuli. A leave-one-participant-out approach was adopted. Specifically, for the 20 participants in each dataset, the data from 19 were used to train a SVM classifier, which was then tested on the remaining participant to obtain decoding accuracy. This process was repeated 20 times and the 20 decoding accuracies were averaged to yield the group-level average decoding accuracy. The reason we took the across-subject decoding approach rather than the more conventional within-subject approach is because for training CNN models (see below), larger amount of data were required, which means that the within-subject analysis was not possible for CNN analysis and the across-subject approach has to be used to increase the size of training data. Although the SVM approach has been mainly applied in a within-subject fashion, for an equitable comparison, we applied the SVM decoding in this study using the same across-subject decoding as the CNN.

In addition to decoding accuracy, the heatmap is another critical aspect of the SVM technique, which can be used to attribute functional significance to voxels inside an ROI. To compute the SVM heatmap inside an ROI, we applied the transformation proposed by Haufe et al. to the weight vectors computed from the SVM (Haufe et al., 2014). The absolute value of the resulting weight in each voxel measures the contribution of the voxel to the decoding performance (Lee et al., 2010; Mourao-Miranda et al., 2005).

### MVPA: CNN

Widely used CNNs such as ResNet (He et al., 2016) take two dimensional images, such as those from the ImageNet (Deng et al., 2009) or Cifar10 (Krizhevsky & Hinton, 2009), as input and produce recognition of the object in the image as output. But fMRI data from the brain are three dimensional. We modified all the convolutional filters to 3D in the original ResNet18 structure without changing the kernel size; see **Figure 3A**. In addition, to make an equitable comparison with the SVM approach, we modified the kernel size of the first convolutional layer, changing it from the original kernel size of 7 to the kernel size of 1. The detailed filter and dimension size was listed in **Table 1**. The same leave-one-participant-out approach as in SVM analysis above was used here for both datasets.

**Figure 3:**
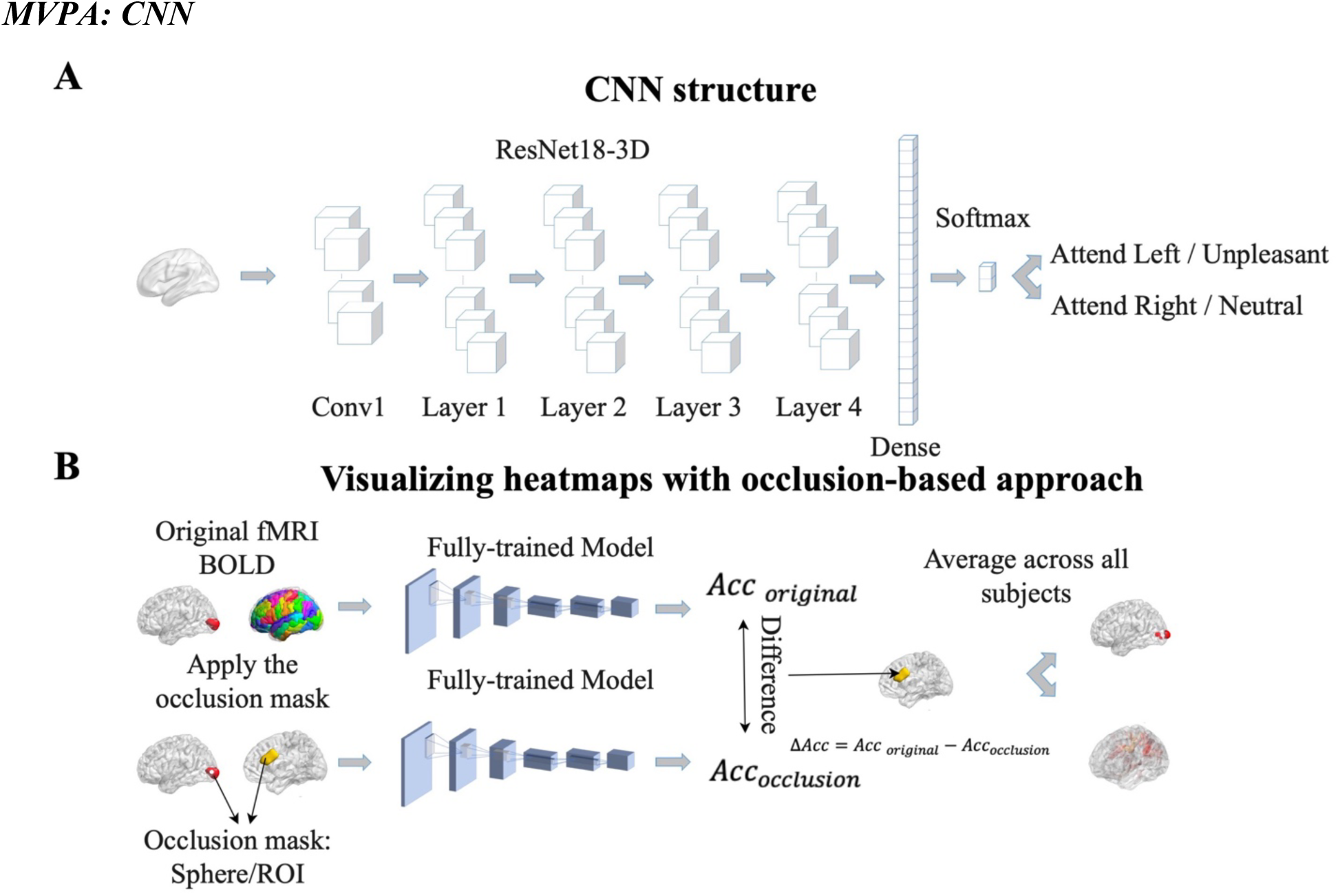
Pipeline for Convolutional Neural Network (CNN) analysis. (A) CNN model was trained to predict attention control or emotional processing from fMRI data. (B) An occlusion approach was carried out to evaluate the contribution of different brain regions or voxels to CNN model classification performance. The detailed filter size and feature map dimension are listed in Table 1. Acc: accuracy.

**Table 1:** Detailed structure of the proposed ResNet-18. F is the number of feature channels, and N is the number of blocks in each layer.

| Model | Block | Conv1 | Layer1 |  | Layer2 |  | Layer3 |  | Layer4 |  | Average Pooling SoftMax |
| --- | --- | --- | --- | --- | --- | --- | --- | --- | --- | --- | --- |
|  |  |  | F | N | F | N | F | N | F | N |  |
| ResNet-18 | ResNet | Conv,1x1x1, 64, stride 2, padding 3 | 64 | 2 | 128 | 2 | 256 | 2 | 512 | 2 |  |

For heatmaps, we applied an occlusion approach. Specifically, for V1 decoding, after a CNN model was trained and the decoding accuracy obtained, we removed a sphere with a radius of 6mm (2 voxels) around each voxel (i.e., change the activity level in the voxels inside the sphere to 0) and fed the fMRI BOLD into the trained CNN network to obtain decoding accuracy. The decrease in decoding accuracy, which is taken as the weight of the voxel in the heatmap, is considered a measure of the significance of the voxel. For whole-brain decoding, after a CNN model was trained and the decoding accuracy obtained, we removed a ROI by setting the activity level in all the voxels in the ROI to 0 and fed the fMRI BOLD into the trained CNN network to obtain decoding accuracy. The decrease in decoding accuracy is taken as a measure of the weight of the ROI in the heatmap, see **Figure 3B**. The logic here is that the more the decoding accuracy decreased following occlusion, the more significant the voxel or ROI contributed to the decoding performance, and the more highly weighted the voxel or ROI is in the heatmap.

## Results

### Attention Dataset: Decoding Attentional Control in V1

According to the prevailing theory of attention control, following an attention directing cue, top-down signals from frontoparietal attention control areas propagate to visual cortex to bias the sensory neurons to enhance attended information and suppress distraction (Corbetta et al., 2008; Liu et al., 2003; Wang et al., 2016) . To what extent such signals reach the level of the primary visual cortex (V1) is not well established. Using V1 as the ROI, we decoded spatial attention control (cue left vs. cue right) using SVM and CNN, and compared the decoding accuracies. As shown in **Figure 4A**, both SVM and CNN had decoding accuracy significantly above chance level of 50% (p<0.0001), with the CNN decoding accuracy significantly higher than the SVM (p<0.002), suggesting that attention control signals can reach the level of V1 and CNN has stronger abilities to detect these signals. Correlating the decoding accuracies between SVM and CNN, we found that the two decoding accuracies were not significantly correlated (R=0.12, p=0.62), suggesting that these two methods may emphasize different data features to achieve their respective decoding performance, see **Figure 4B**.

**Figure 4:**
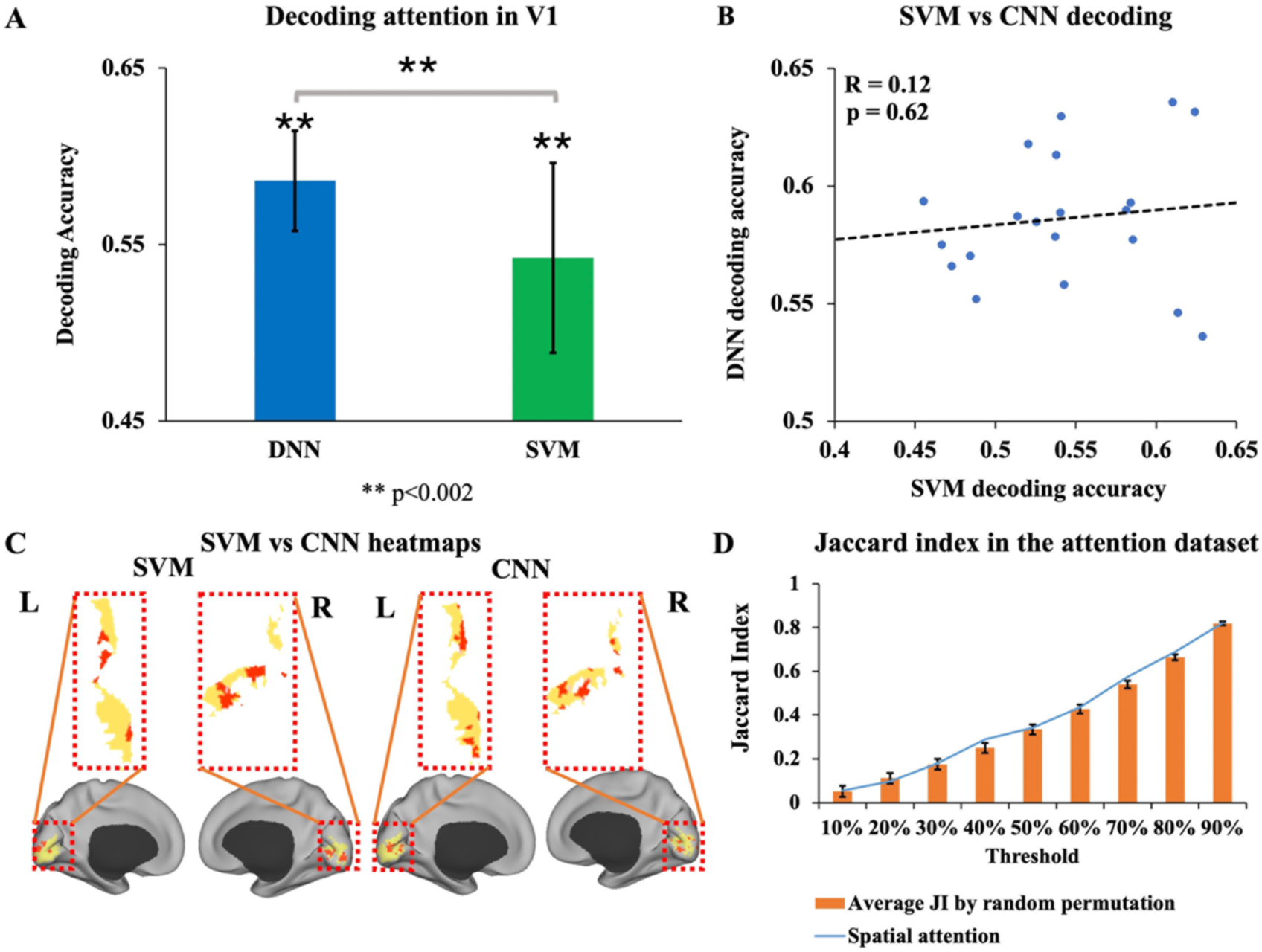
Decoding attention control in V1. (A) SVM versus CNN decoding. Both decoding accuracies are significantly above chance level (p<0.0001), with CNN decoding accuracy higher than SVM decoding accuracy (p<0.002). (B) Correlation between SVM and CNN decoding accuracies. The decoding accuracies from the two methods are not significantly correlated (R=0.12, p=0.62). (C) The heatmaps underlying spatial attention control by SVM and CNN. The two heatmaps have limited overlap for the top 20% of voxels (JI=0.10), suggesting that the two methods emphasize different features for making predictions. (D) The average random permutation JI and the JI from the data at different thresholds from 10% to 90%. None of the thresholds showed significant differences between the random permutation JI and the JI from the data. JI: Jaccard Index. *\*\*p<0.002*

To examine whether different features drove the decoding performance for the two different kinds of MVPA methods (SVM vs CNN), we compared the top 20% voxels (80 voxels) from the heatmap of SVM and that of CNN; **see Figure 4C**. Intuitively the heatmaps look quite different. To quantify the extent of overlap/nonoverlap of two sets of voxels in the two heatmaps, the Jaccard index (JI) was computed, where a JI of 0 or 1 indicates no overlap or total overlap (Levandowsky & Winter, 1971). We found JI=0.10 to be the overlap between the top 80 voxels from the SVM heatmap and that from the CNN map. To better understand the meaning of this number, we randomly chose two sets of 80 voxels and computed the JI. Repeating this procedure 10000 times, the average random permutation JI was found to be 0.11 ± 0.025. Comparing JI=0.10 from the actual data with the JI from random permutation yielded no significant difference (p=0.65). We take this as evidence that SVM and CNN rely on different feature information for making classification of the two attentional states. Extending the percent of voxels included in the overlap analysis to [10%,90%], we found that the JI is an increasing function of the percent threshold, as shown in **Figure 4D**, and at every threshold, the JI from the data is not significantly different from the JI of two sets of randomly chosen voxels (p>0.05, FDR corrected). These additional experimental results confirm that SVM and CNN rely on different voxel features for representing different attentional control states (attend left vs attend right).

### Attention Dataset: Decoding Attentional Control over the Whole Brain

Traditional whole-brain univariate analysis was done first to provide a basis for comparison with the subsequent multivariate analysis. As in previous work, the whole-brain responses evoked by spatial cues (left cue and right cue combined) were analyzed using the GLM method. As depicted in **Figure 5A**, the spatial cues activated the precentral/postcentral cortex which contains the frontal eye field (FEF) and superior parietal cortex which contains the inferior parietal sulcus (IPS)/superior parietal lobule (SPL), which is consistent with previous findings (Giesbrecht et al., 2003; Rajan et al., 2021; Slagter et al., 2007); FEF and IPS/SPL comprise the dorsal attention network (DAN). Additionally, other regions such as the putamen, superior frontal and inferior frontal were also activated, agreeing with our previous study (Meyyappan et al., 2021). A total of 9 regions are activated by the spatial cues (see **Table 2**). However, when the left cue is contrasted again the right cue or vice versa, no regions appeared in the activation map, suggesting that the regions that encode the specific attended information are not discovered by the univariate analysis; see **Figure 5B** and **5C**.

**Figure 5:**
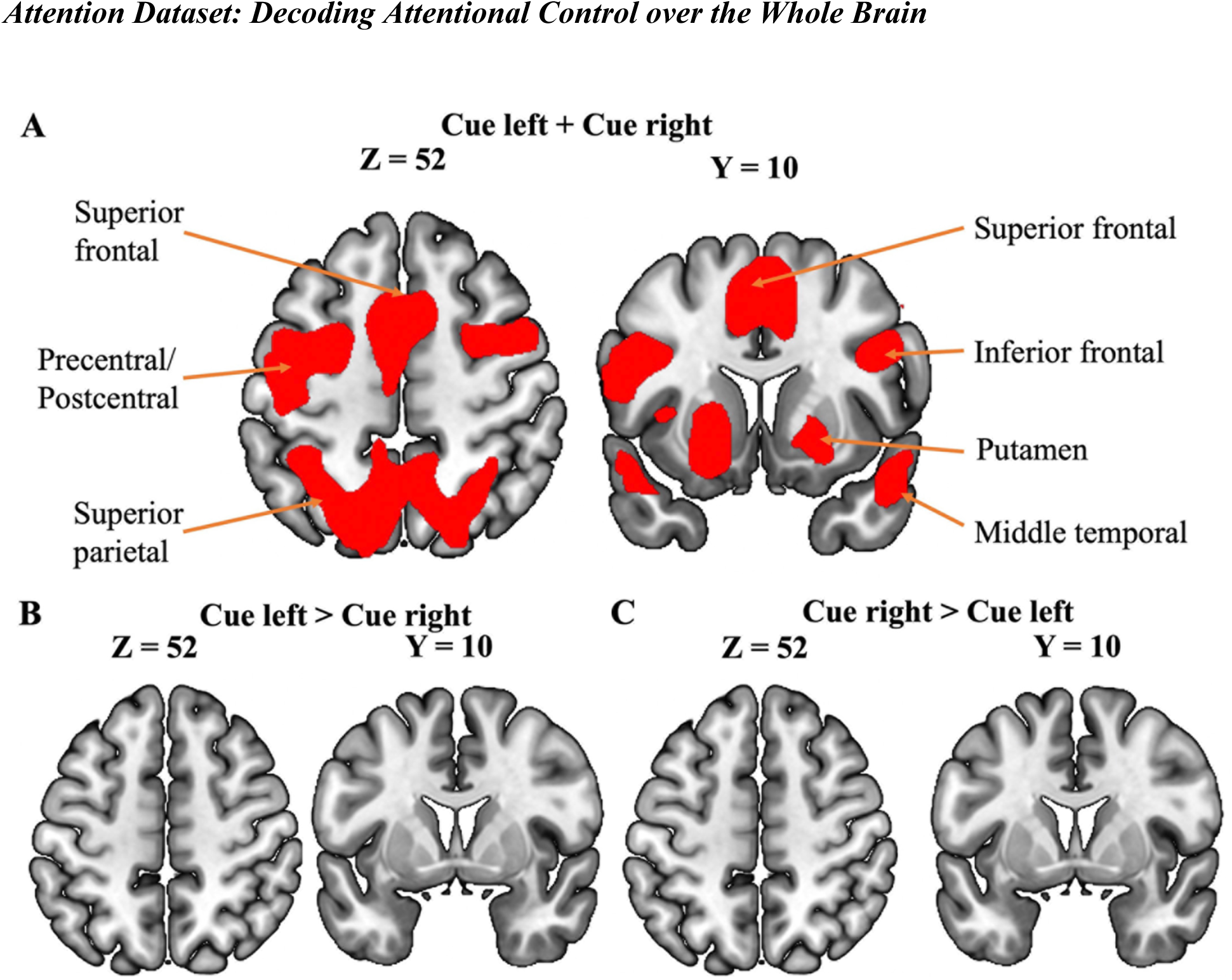
Univariate analysis of attention control. (A) Cue-evoked activation map (*p* < 0.05, FDR) combining cue left and cue right. Several regions, including precentral/postcentral, superior parietal, inferior frontal, and putamen, were activated. (B) Cue left > cue right activation map (*p* < 0.05, FDR). No regions appeared in the map. (C) Cue right > cue left activation map (*p*< 0.05, FDR). No regions appeared in the map.

**Table 2:** Important regions underlying attention control identified by univariate and MVPA analysis.

|  | Cue left + Cue right | Cue left vs Cue right | SVM | CNN |
| --- | --- | --- | --- | --- |
| Middle temporal | √ |  |  |  |
| Bankssts | √ |  |  |  |
| Lateral orbito frontal | √ |  |  |  |
| Precentral | √ |  |  | √ |
| Lateral occipital | √ |  | √ | √ |
| Fusiform | √ |  | √ | √ |
| Insula | √ |  | √ |  |
| Rostral middle frontal | √ |  |  | √ |
| Medial orbito frontal | √ |  |  |  |
| Superior parietal |  |  | √ | √ |
| Parsorbitalis |  |  | √ |  |
| Pericalcarine |  |  | √ |  |
| Parstriangularis |  |  | √ |  |
| Lingual |  |  | √ |  |
| Cuneus |  |  | √ |  |
| Inferior temporal |  |  |  | √ |
| Thalamus |  |  |  | √ |
| Inferior parietal |  |  |  | √ |
| Superior temporal |  |  |  | √ |

In contrast, both the MVPA methods SVM and CNN were able to decode the two attention control conditions (cue left vs. cue right) in the whole-brain analysis (p<0.0001), as shown in **Figure 6A**, with the CNN decoding accuracy significantly higher than SVM decoding accuracy (p<0.05). Unlike in V1, over the whole brain, the decoding accuracies between SVM and CNN were significantly correlated (R = 0.77, p<0.0001), suggesting the at the whole brain level, the two decoding methods may rely more on common input features than totally different input features; see **Figure 6B**.

**Figure 6:**
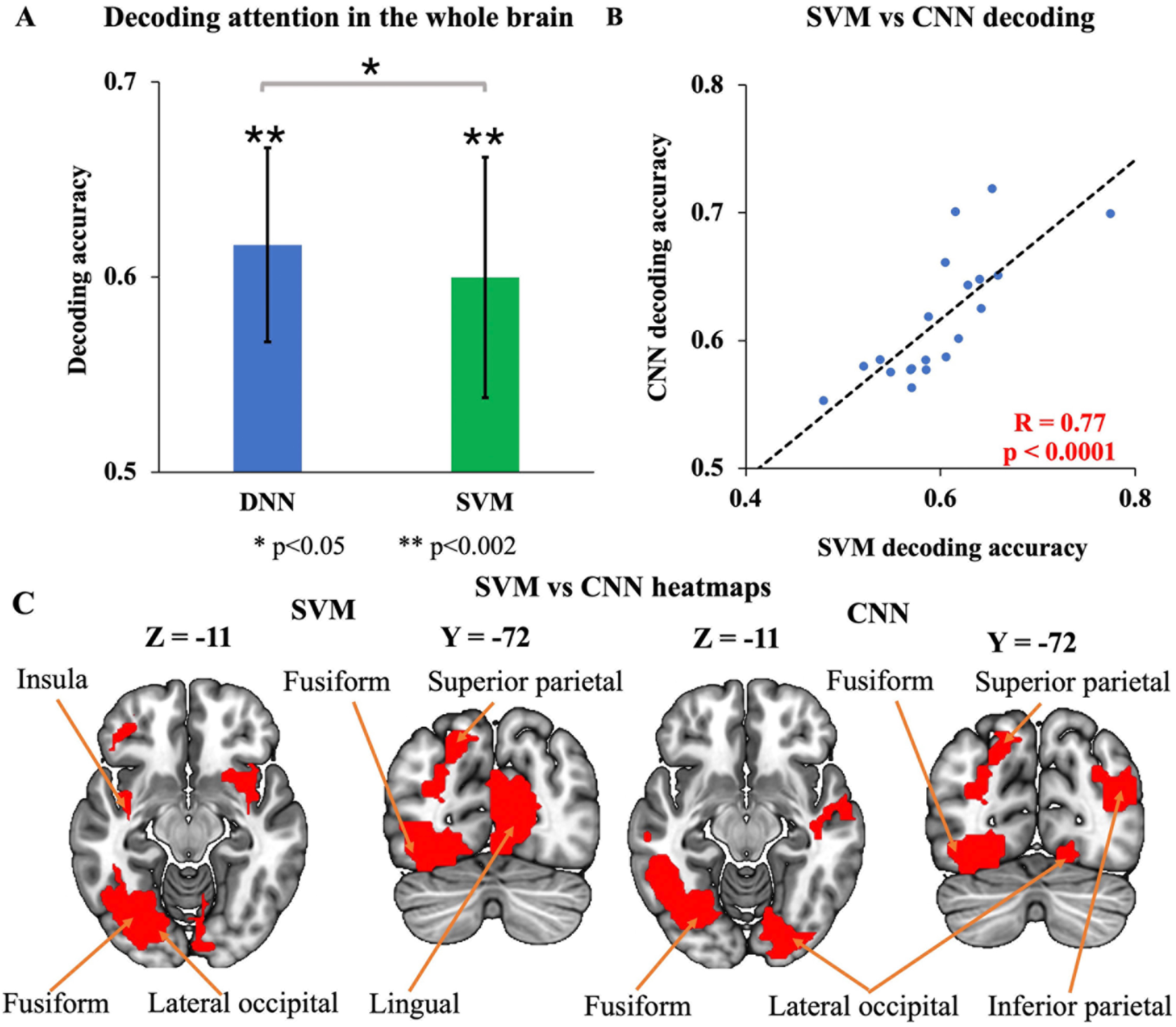
Decoding attention control over the whole brain. (A) fMRI spatial attention decoding accuracy over the whole brain by SVM and CNN. Both decoding accuracies are significantly above chance level (p<0.0001), with CNN decoding accuracy higher than SVM decoding accuracy (p<0.05). (B) Correlation between SVM and CNN decoding accuracies for spatial attention over the whole brain. The decoding accuracies from the two methods are significantly correlated (R=0.77, p<0.0001). (C) The heatmaps underlying spatial attention control. The two heatmaps have several overlapping ROIs, including the fusiform, lateral occipital, and superior parietal cortex. \*\**p<0.002* \**p<0.05*

To further examine the features that drove the classification performance, instead of focusing on voxels as in the case of V1, we focused on the regions of interest for ease of interpretation and comparison with univariate analysis. For SVM and CNN, we presented the top 9 regions in the heatmap (which was the same number of regions as those identified in the univariate analysis) from each method, as shown **in Figure 6C**. The top 9 regions identified by the univariate and each of the two MVPA methods are listed in **Table 2**. There were very few overlapping ROIs among the methods. For example, only lateral occipital, fusiform and superior parietal regions was identified by both SVM and CNN. The lateral occipital, fusiform, and insula appeared in both univariate and SVM analyses, and the precentral, lateral occipital, fusiform and the rostral middle frontal were the areas appearing in both univariate and CNN analyses. In summary, each of the three methods provided insights that are largely not contained in the other methods, suggesting that combining these methods might be a way to obtain a more comprehensive understanding of spatial attention control.

### Emotion Dataset: Decoding Emotion Processing in V1

Conventionally, the primary visual cortex (V1) is thought to mainly play the role of extracting basic visual features from sensory input and send them to higher order visual areas for further processing. Our recent work where decoding is applied in a within-subject manner found affect-specific neural representations in V1 when participants viewed natural images containing varying degrees of affective content (Bo et al., 2021). Here, for the same dataset, as shown in **Figure 7A** and **7B**, both SVM and CNN, when applied in a across-subject fashion, exhibited decoding accuracies between unpleasant and neutral images that were significantly higher than chance level of 50% (p<0.0001), with CNN decoding accuracy significantly higher than that of SVM (p<0.002), further confirming that V1’s role is not limited to extracting visual features and it forms an integral part of the neural network that represents the affective significance of natural images.

**Figure 7:**
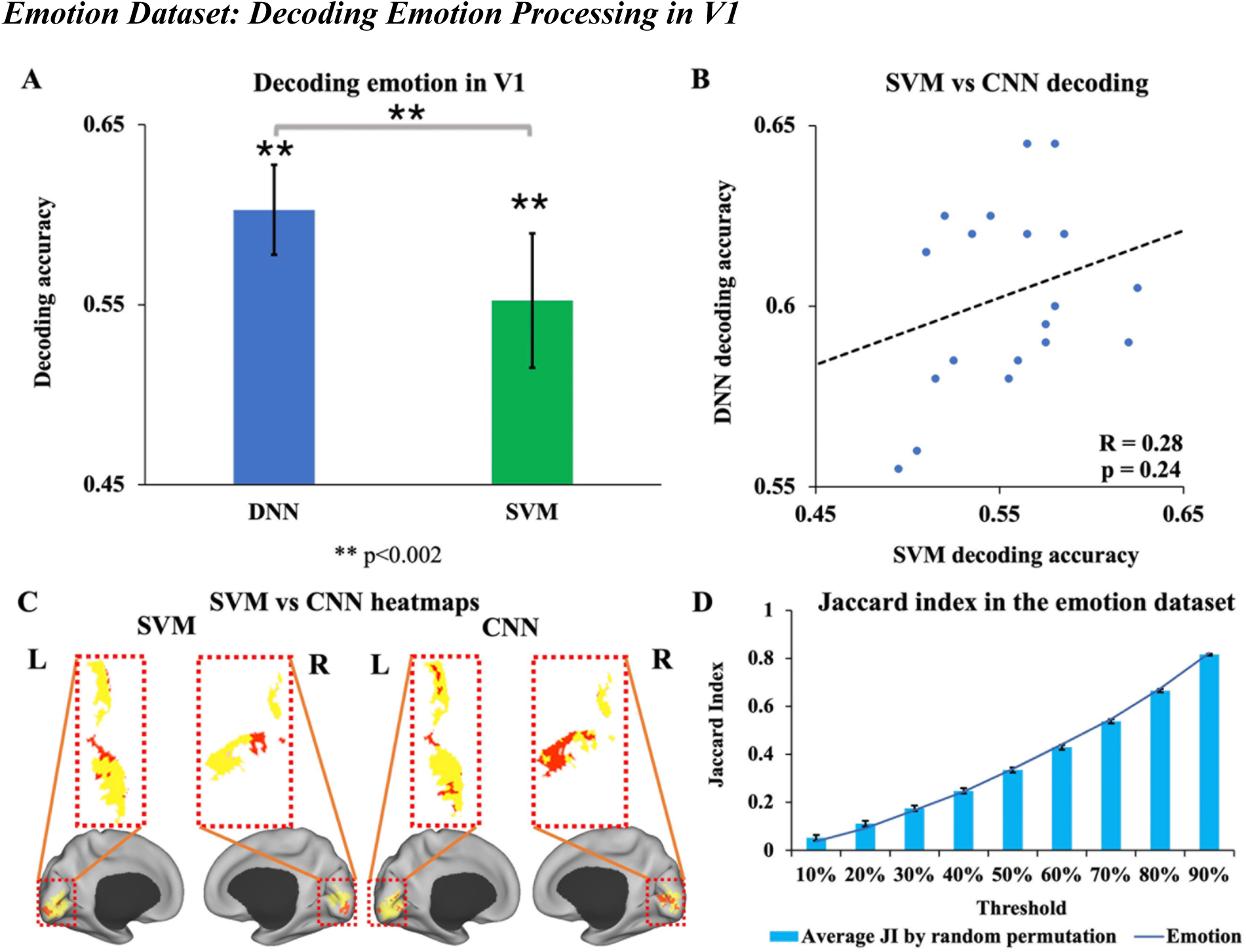
Decoding emotion processing in V1. (A) Decoding unpleasant vs neutral images in V1 by SVM and CNN. Both decoding accuracies are significantly above chance level (p<0.0001). CNN decoding accuracy is higher than SVM decoding accuracy (p<0.002). (B) Correlation between SVM and CNN decoding accuracies. The decoding accuracies from the two methods are not significantly correlated (R=0.28, p=0.24). (C) The heatmaps underlying decoding by SVM and CNN. The two heatmaps have limited overlap for the top 20% of voxels (JI=0.09), suggesting that the two methods may emphasize different features for making predictions. (D) The average random permutation JI and the JI from the data at different thresholds from 10% to 90%. None of the thresholds showed significant differences between the random permutation JI and the JI from the data. JI: Jaccard Index. *\*\*p<0.002*

To test whether the same or different features drove the SVM vs CNN classification performance, we compared the heatmaps from the two methods. The top 20% of voxels (84 voxels) from the SVM heatmap and CNN heatmap were shown in **Figure 7C**. Intuitively, there is little overlap between the two sets of voxels; quantitatively, the JI=0.09. Choosing two random sets of 84 voxels and computing the JI 10000 times, the average JI was found to be 0.11 ± 0.024. The JI of 0.09, which is not significantly different from the overlap between two sets of random voxels (R=0.28, p=0.24). Extending the voxel selection threshold from 10% to 90% and computing the JI index for each threshold, we found that the JI value increased as percentage threshold was increased, as would be expected, but for every threshold, the JI from the data is not significantly different from the overlap of two random sets of voxels (p>0.05, FDR corrected); see **Figure 7D**. These results suggest that the features driving the prediction of SVM and CNN differ.

### Emotion Dataset: Decoding Emotional Processing over the Whole Brain

We first conducted a whole-brain univariate analysis by contrasting unpleasant images against neutral images. In comparison to neutral pictures, unpleasant pictures evoked activations in the precentral, insula, putamen, amygdala, and inferior temporal cortices, as shown in **Figure 8A**. On the other hand, relative to unpleasant pictures, neutral pictures activated the lateral orbitofrontal, superior frontal, middle temporal and parahippocampal regions, as illustrated in **Figure 8B**. The univariate activation map in **Figure 8** was thresholded at p<0.05 FDR. The regions in **Figures 8A** and **8B** are given in **Table 3**. Prior empirical investigations have provided strong evidence for the pivotal roles of the amygdala, insula and parahippocampal in the processing of pain and emotion. Our univariate analysis results are thus consistent with the reports in the literature.

**Figure 8:**
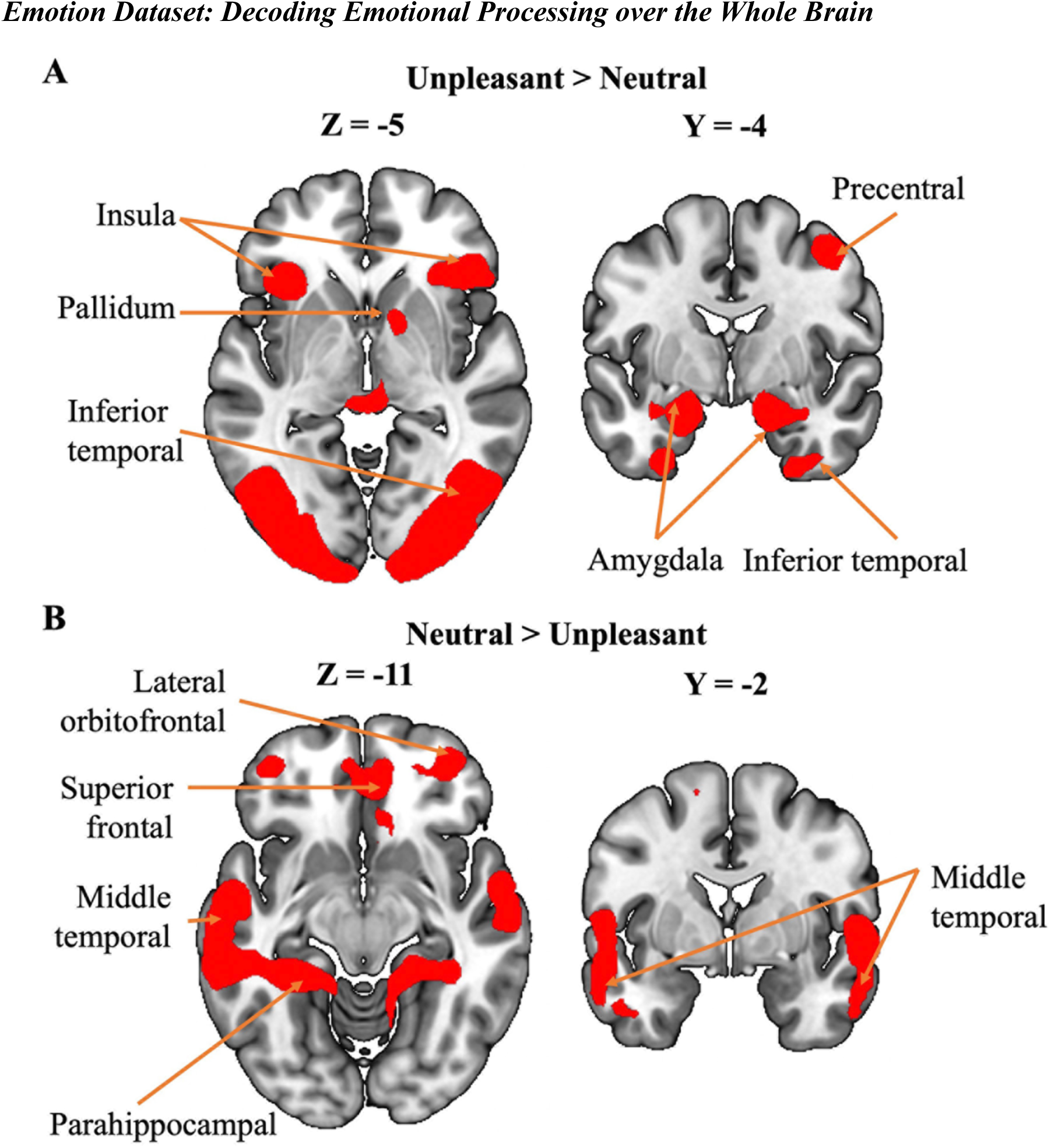
Activation map (p < 0.05, FDR) contrasting (A) unpleasant > neutral (B) neutral > unpleasant pictures.

**Table 3:** Important regions underlying emotion processing identified by univariate and MVPA analysis.

|  | Unpleasant>Neutral | Neutral>Unpleasant | SVM | CNN |
| --- | --- | --- | --- | --- |
| Inferior temporal | √ |  | √ |  |
| Lateral orbito frontal | √ |  |  |  |
| Amygdala | √ |  |  |  |
| Superior frontal | √ |  |  |  |
| Dorsal Acc | √ |  |  |  |
| Brainstem | √ |  |  | √ |
| Pallidum | √ |  |  |  |
| Cuneus |  | √ | √ |  |
| Caudal middle frontal |  | √ |  |  |
| Rostral middle frontal |  | √ |  | √ |
| Rostral Acc |  | √ |  |  |
| Fusiform |  |  | √ | √ |
| Pericalcarine |  |  | √ |  |
| Lingual |  |  | √ |  |
| Lateral occipital |  |  | √ | √ |
| Supramarginal |  |  | √ |  |
| Inferior parietal |  |  | √ | √ |
| Precunues |  |  | √ | √ |
| Posterior cingulate |  |  | √ |  |
| Superior parietal |  |  | √ |  |
| Middle temporal |  |  |  | √ |
| Pars opercularis |  |  |  | √ |
| Superior temporal |  |  |  | √ |
| Medial orbito frontal |  |  |  | √ |
| Parsorbitalis |  |  |  | √ |

Both MVPA methods SVM and CNN achieved decoding accuracies that were significantly higher than chance level (p<0.0001), with the CNN decoding accuracy significantly higher than that of SVM (p<0.05); see **Figure 9A**. No significant correlations were found between the decoding accuracies of SVM and CNN in **Figure 9B** (R=0.07, p=0.78). To what extent this finding implies that the two methods relied on different features to perform classifications was examined next using the heatmap.

**Figure 9:**
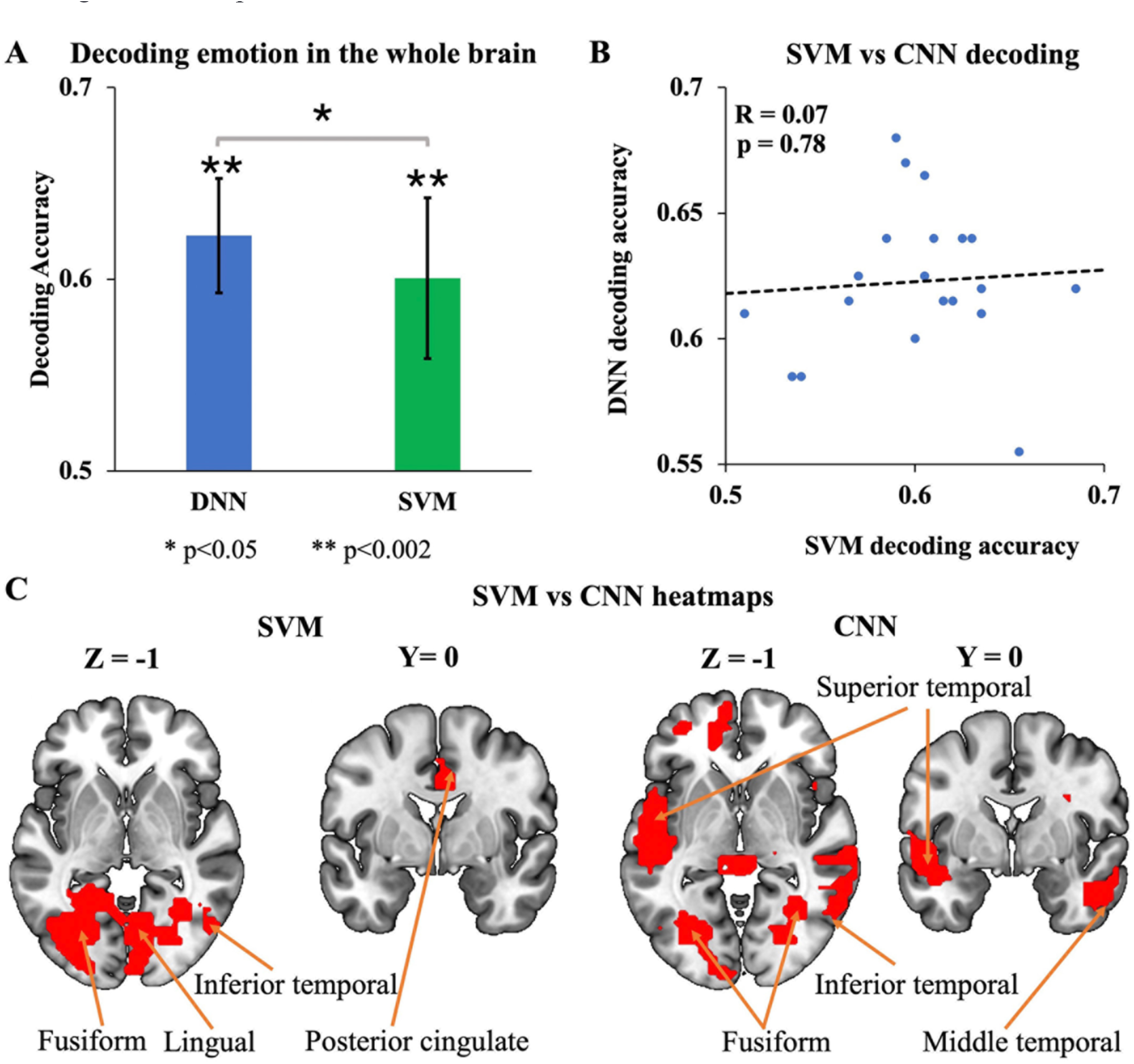
Decoding unpleasant vs neutral over the whole brain. (A) fMRI decoding accuracy in the whole brain by SVM and CNN. Both decoding accuracies are significantly above chance level (p<0.0001). CNN achieved significantly higher decoding accuracy compared to SVM (p<0.05). (B) Correlation between SVM and CNN decoding accuracies. The decoding accuracies from the two methods are not significantly correlated (p=0.78). (C) The heatmaps underlying the decoding between unpleasant vs. neutral over the whole brain. Both heatmaps include the fusiform, inferior temporal, lateral occipital, and precuneus as essential brain regions. \*\**p<0.002* \**p<0.05*

To identify the essential ROIs driving the prediction performance in both SVM and CNN, we selected the top 12 ROIs (same as the number of ROI identified in the univariate analysis) based on the Lausanne atlas as critical ROIs for generating the heatmaps, as shown in **Figure 9C**. Regions including fusiform, lateral occipital, inferior temporal and precuneus were detected simultaneously by two approaches. See **Table 3.**

## Discussion

MVPA methods are now widely used in neuroimaging. SVM has been the mainstay in MVPA analysis of fMRI data since the early 2000s. In more recent years DNNs especially CNNs have been added to the MVPA’s toolchest and are gaining importance due to its ability to take into account of nonlinear relationships between neural patterns and cognitive functions. To date no systematic comparisons between these two classes of methods have been carried out. Using two well-characterized fMRI datasets, we compared the decoding performance and heatmaps derived from SVM and CNN, and reported the following results. First, both SVM and CNN are able to achieve above chance level decoding accuracies for attention control and emotion processing in both the primary visual cortex and the whole brain, and the CNN decoding accuracies were consistently higher than that of the SVM. Second, the SVM and CNN decoding accuracies are generally not correlated, and the heatmaps derived by SVM and CNN, which emphasize the importance of certain voxels or brain regions over others in decoding performance, were shown to be not significantly overlapping, suggesting that SVM and CNN rely on different brain activity patterns to predict cognitive conditions.

### Analysis in V1

The primary visual cortex (V1) is the first stage of the cortical visual processing system (Hubel & Wiesel, 1977). As such its role is thought to be limited to extracting visual features from input and passing them up the visual hierarchy for further processing. For both datasets, our results demonstrate that V1’s role goes beyond simple feature extraction. In the case of visual spatial attention, while attentional modulation of neuronal responses has been reported in many extrastriate cortical areas such as V2, V4, temporal-occipital area (TEO), and middle temporal area (MT), evidence of attentional modulation in V1 has been relatively scarce (Pessoa et al., 2003), even though it is well known that V1 receives feedback connections from higher-level visual areas, allowing its activity to be modulated by top-down attentional control signals (Juan & Walsh, 2003). In the present study, both SVM and CNN achieved above-chance decoding accuracies between two preparatory attention conditions (attend left vs attend right) using only the voxels extracted from V1, demonstrating a role of V1 in attention control. Furthermore, owing to the across-subject decoding approach adopted here, the patterns of the attention control signals are found to be shared by the participants (Martinez et al., 1999). In the case of visual emotion processing, the role of V1 (primary visual cortex) is not well understood, and findings from different studies have been inconsistent. Generally, univariate BOLD analyses have not found differential activations by differently valenced emotional images (e.g., unpleasant vs neutral) in early visual areas such as V1 (Sabatinelli et al., 2009). Using a within-subject decoding approach, our previous analysis of the same emotion dataset shows that there are affect-specific neural representations in the retinotopic visual cortex, which includes V1 (Bo et al., 2021). In the present study, applying an across-subject approach using two different decoding methods, we again found evidence to support the notion that the primary visual cortex V1 plays a role in the representation of the emotional significance of natural images and these representations are shared by participants. It is worth noting that for both datasets, the heatmaps derived from SVM and CNN are largely nonoverlapping, suggesting that these two classes of methods may rely on different features for decoding. The nature of these features remains not clear and should be the subject of future studies.

### Analysis in whole-brain

At the whole brain level, we first performed the traditional univariate analysis to provide a departure point for the comparison of the two MVPA methods. For the attention dataset, when attend left cue and attend right cue were combined, the frontal eye field and the IPS/SPL were found to be activated. These regions are part of the dorsal attention network (DAN) whose role in the control of spatial attention, feature attention, and object attention is well established (Corbetta et al., 2005; Giesbrecht et al., 2003; Morishima et al., 2009; Slagter et al., 2007). Additional areas activated by the combined cues include inferior frontal cortex, which are associated with cognitive control and response inhibition, both being integral parts of attention (Forstmann et al., 2008). When we contrasted attend left cue vs attend right cue, no brain activations were found, suggesting that the univariate analysis is not able to discover the signals that are specific to the attended information (e.g., attend left vs attend right). In contrast, both SVM and CNN were able to decode attend left vs attend right using the whole brain as the ROI. In particular, these MVPA analyses were able to unveil brain regions whose involvement in attention control was not uncovered by the conventional univariate analysis, including the inferior parietal cortex and inferior temporal cortex. Recent studies utilizing imaging techniques and lesion investigations have provided insights into the functional properties of inferior parietal regions and found that they contribute to the maintenance of attention, detection of salient events within a sequence, and exertion of control over attentional mechanisms (Husain & Nachev, 2007). Moreover, event-related fMRI investigations have identified a network of cortical areas, including the inferior parietal cortex, that play a role in top-down attentional control (Hopfinger et al., 2000) and attentional shifting (Corbetta et al., 1993). In terms of the univariate activation map and the heat maps derived from the two MVPA methods, similar to V1, there is limited overlap among the regions identified by each of the three methodologies (i.e., univariate, SVM, and CNN) as being important for spatial attention control, suggesting that combining these methods may afford us the ability to more comprehensively identify important neural substrate of cognitive functions. One observation worth noting is that the DAN did not appear in the heatmap of either of the two MVPA methods. Our previous decoding analysis in DAN, using a within-subject approach (Rajan et al., 2021), demonstrated the existence of a microstructure for attention control in DAN, consistent with an extensive literature (Giesbrecht et al., 2003; Morishima et al., 2009). The findings reported here may be taken to imply that the microstructure of attention control exhibits a high degree of individual variability, which renders the across-subject decoding approach not effective in revealing its involvement in attentional control.

For the emotion dataset, the univariate analysis revealed activations in multiple brain areas when contrasting unpleasant images against neutral images, including the insula, precentral gyrus, amygdala, fusiform gyrus, superior frontal gyrus, and middle temporal gyrus, as reported in our previous study (Bo et al., 2021). The amygdala and insula are widely recognized as key brain regions involved in processing emotional information, particularly negative emotions (Kesler et al., 2001). The superior frontal region is implicated in the cognitive ability to infer the mental states of others and other social functions where emotion processing is critical (Mak et al., 2009; Völlm et al., 2006). Notably, the retinotopic visual cortex is not engaged in representing the emotional significance of visual processing according to the univariate analysis, contrary to the theoretical argument. Specifically, the signal reentry hypothesis suggested that subcortical structures, such as the amygdala, may send feedback signals into the visual cortex upon receiving initial sensory input, thereby enhancing the processing of emotionally salient visual stimuli (Keil et al., 2009; Lang & Bradley, 2010; Sabatinelli et al., 2009). In both SVM and CNN analysis, the visual cortex was found contributing to the decoding performance, in agreement with our previous study using a within-subject decoding approach (Bo et al., 2021) and suggesting that the neural representations of the emotional significance of natural images are shared across participants. Furthermore, the involvement of the precuneus in representing emotion was found by both SVM and CNN, but was not revealed by univariate analysis. The precuneus region, along with activity in the right anterior insular cortex and ventromedial prefrontal cortex (VMPFC), has been linked to the evaluation of emotional states (Terasawa et al., 2013). Critchley et al. (Critchley et al., 2001) provides evidence that the precuneus plays a critical role in transforming interoceptive information into subjective emotions.

Summarizing the foregoing we note that univariate analysis and MVPA employ different approaches and can offer complementary information. Univariate analysis typically identifies regions that exhibit statistically significant differences in activity levels, while MVPA provides insights into how patterns of activity across multiple voxels or across multiple regions are related to specific experimental conditions or stimuli. Therefore, combining both univariate and MVPA approaches can lead to a more comprehensive understanding of the neural mechanisms underlying attention processes and emotional perception in fMRI research.

### Methodological Considerations and Limitation

Two methodological issues are worth considering. First, the findings of the current study suggest that CNN performed better than SVM in all decoding tasks. This advantage of CNN can be attributed to its superior learning abilities relative to SVM (Khodatars et al., 2021); SVM can be considered as having a single-layer architecture and is thus less computationally capable compared to a multi-layered system such as the CNN (Kim et al., 2019). However, CNN, having multiple layers of computational units (neurons), has a large number of parameters; it is known that as the number of parameters is increased, overfitting is more likely to occur. We applied the traditional cross-validation approach, batch normalization, and dropout in the current study to avoid overfitting (Ioffe & Szegedy, 2015; Srivastava et al., 2014). In future research, additional methods such as L1 and L2 norm regularizations, as well as data augmentation techniques, could be considered to address the issue of relatively small sample sizes in neuroimaging datasets. Second, we utilized an across-subject decoding approach rather than the more traditional within-subject decoding approach. In the within-subject decoding approach, patterns of neural activity are collected for each individual during a specific experimental condition or task, and then subjected to decoding or classification analysis. The across-subject approach, which is adopted here to cope with the fact that training CNN models requires larger amount of data than can be expected from a single subject in typical neuroimaging studies, involves pooling data from multiple participants performing the same experiment, and then performing decoding or classification of the patterns; the decodable patterns in this case are patterns that are common across the group. The within-subject and across-subject decoding approaches each have their advantages and limitations. Within-subject decoding allows for investigating individual differences and can provide insights into the neural mechanisms that are specific to each individual. It is less affected by inter-individual variability and can potentially capture more fine-grained individual-specific information. The use of an across-subject approach in fMRI research also has advantages. For example, pooling data from multiple subjects can increase the statistical power of the analysis, making it more likely to detect subtle or complex effects that may not be apparent in individual subjects due to variability or noise in the data.

